# Supramolecular Interactions of Teixobactin Analogues in the Crystal State

**DOI:** 10.1101/2023.10.30.564786

**Authors:** Hyunjun Yang, Adam G. Kreutzer, James S. Nowick

## Abstract

Teixobactin is a potent peptide antibiotic against Gram-positive bacteria that binds to lipid II and related peptidoglycan precursors and disrupts the cell membrane. This paper presents the X-ray crystallographic structure of the *N*-methylated teixobactin analogue *N*-Me-D-Gln_4_,Lys_10_-teixobactin (1). *N*-Methylation at position 4 prevents uncontrolled aggregation and enables the crystallization of the teixobactin analogue. Lysine at position 10 replaces the non-proteinogenic amino acid *allo*-enduracididine, which is not commercially available. Crystallization from aqueous solution with MgCl_2_ and PEG3350 afforded crystals suitable for X-ray crystallography. The crystallographic phases were solved using SAD phasing on data sets collected at 2.0663 Å. Molecular replacement then enabled the determination of the structure at 1.50 Å resolution using a data set collected at 0.9997Å (PDB 8U78). Eight peptide molecules comprise the asymmetric unit, with each peptide molecule binding a chloride anion through hydrogen bonding with the amide NH group of residues 7, 8, 10, and 11. The peptide molecules form hydrogen-bonded antiparallel β-sheet dimers in the crystal lattice, with residues 1–3 comprising the dimerization interface. The dimers further assemble end-to-end in the crystal lattice. The β-sheet dimers are amphiphilic, with the side chains of the hydrophobic residues on one surface and the side chains of the hydrophilic residues on the other surface. The dimers pack in the lattice through hydrophobic interactions between the hydrophobic surfaces. The crystal structure of teixobactin analogue 1 recapitulates several aspects of the interaction of teixobactin with the cell membrane of Grampositive bacteria, including anion binding, supramolecular assembly through β-sheet formation, and hydrophobic interactions.

## INTRODUCTION

Teixobactin is a peptide antibiotic that exhibits remarkable efficacy against Gram-positive bacteria, including methicillin-resistant *Staphylococcus aureus*, drug-resistant *Streptococcus pneumonia*, and vancomycin-resistant *Enterococci*. ^1^ Teixobactin’s mode of action involves binding to lipid II and related peptidoglycan precursors and disrupting the bacterial cell membrane. Teixobactin is an undecapeptide consisting of an *N*-terminal linear tail encompassing residues 1 to 7 and a *C*-terminal macrocyclic ring spanning residues 8 to 11 (**Figure 1**). The linear tail consists of *N*-Me-D-Phe_1_, Ile_2_, Ser_3_, D-Gln_4_, D-*allo*-Ile_5_, Ile_6_, and Ser_7_, and the macrocyclic ring consists of D-Thr_8_, Ala_9_, the cyclic arginine analogue *allo*-End_10_, and Ile_11_. The *C*-terminus of Ile_11_ and the hydroxy group of D-Thr_8_ form an ester bond, creating 13-membered lactone ring.

**Figure 1.**
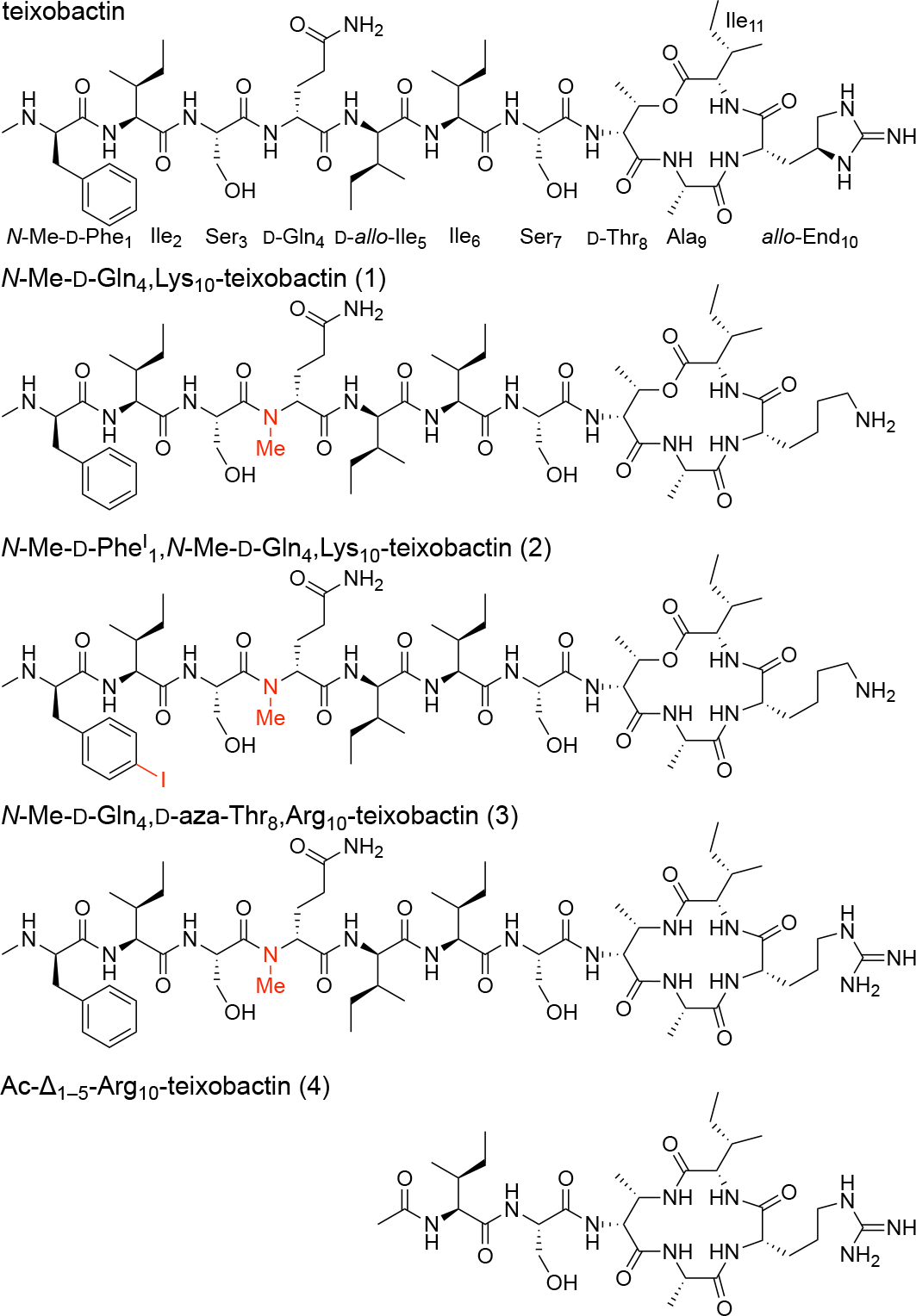
Chemical structures of teixobactin, *N*-methylated D-Gln_4_ analogues of teixobactin, and a truncated teixobactin analogue.

Supramolecular assembly is central to the mechanism of action of.^2,3^ Two teixobactin molecules come together to form an antiparallel β-sheet dimer which binds the pyrophosphate group of lipid II. The *N*-terminal tails comprise the dimerization interface and adopt a conformation in which the residues *N*-Me-D-Phe_1_, Ile_2_, D-*allo*-Ile_5_, and Ile_6_ form a hydrophobic surface that interacts with the bacterial cell membrane. Further supramolecular assembly leads to clustering of lipid II and lysis of Gram-positive bacteria.

Our laboratory has pioneered the use of X-ray crystallography to gain insights into the structure and supramolecular interactions of teixobactin. Although teixobactin itself is highly prone to amyloid-like aggregation and cannot be crystallized, our laboratory has found that *N*-methylation at D-Gln_4_ blocks aggregation and permits crystallization. The resulting *N*-methylated teixobactin analogues exhibit little or no antibiotic activity. We previously observed that teixobactin analogue 2 undergoes supramolecular assembly to form antiparallel β-sheet dimers that bind sulfate anions and further form fibril-like assemblies (PDB 6E00). ^4^ We have also observed antiparallel β-sheet dimers in the X-ray crystallographic structure of teixobactin analogue 3, and we have observed that these dimers bind chloride anions (PDB 6PSL).^5^ We have further observed chloride anion binding in the X-ray crystallographic structure of truncated teixobactin analogue 4 (CCDC 1523518).^6^ In the current study, we report the X-ray crystallographic structure of teixobactin analogue 1 and describe its supramolecular interactions in the crystal state.

## RESULTS

The X-ray crystallographic structure of teixobactin analogue 1 reveals eight peptide molecules in the asymmetric unit, with each peptide molecule binding a chloride anion (**Figure 2**). The eight peptide molecules adopt similar conformations, with a backbone RMSD of 0.58 Å. Each peptide molecule forms an amphiphilic surface in which the side chains of *N*-Me-D-Phe_1_, Ile_2_, D-*allo*-Ile_5_, Ile_6_, Ile_11_, and the β-methyl of D-Thr_8_ create a hydrophobic surface and the side chains of residues Ser_3_, *N*-Me-D-Gln_4_, Ser_7_, and Lys_10_ create a hydrophilic surface. The carbonyl groups of the macrocycle align, and the amide NH groups of Ser_7_, D-Thr_8_, Lys_10_, and Ile_11_ hydrogen bond to chloride anion. The oxygen atom comprising the ester linkage also faces toward the chloride ion. The amide NH group of Ala_9_ hydrogen bonds to the hydroxy group of Ser_7_.

**Figure 2.**
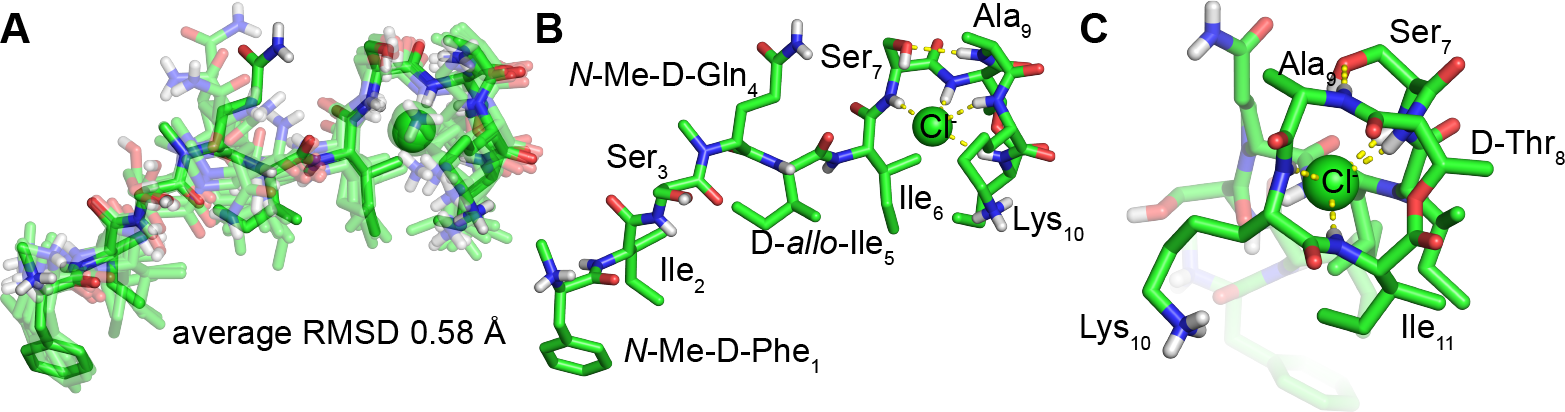
X-ray crystallographic structure of teixobactin analogue 1 (PDB 8U78). (A) Overlay of the eight peptide molecules binding chloride anions in the asymmetric unit. (B) Side view and (C) end view of a representative peptide molecule.

The peptide molecules form hydrogen-bonded antiparallel β-sheet dimers in the crystal lattice (**Figure 3**). In each dimer, residues 1–3 come together through hydrogen bonding interactions to create the antiparallel β-sheet structure. The *N*-methyl groups of *N*-Me-D-Gln_4_ point outward from the hydrogen-bonding interface. The dimer is amphiphilic, with the top surface shown in **Figure 3** being hydrophilic and the bottom surface being hydrophobic.

**Figure 3.**
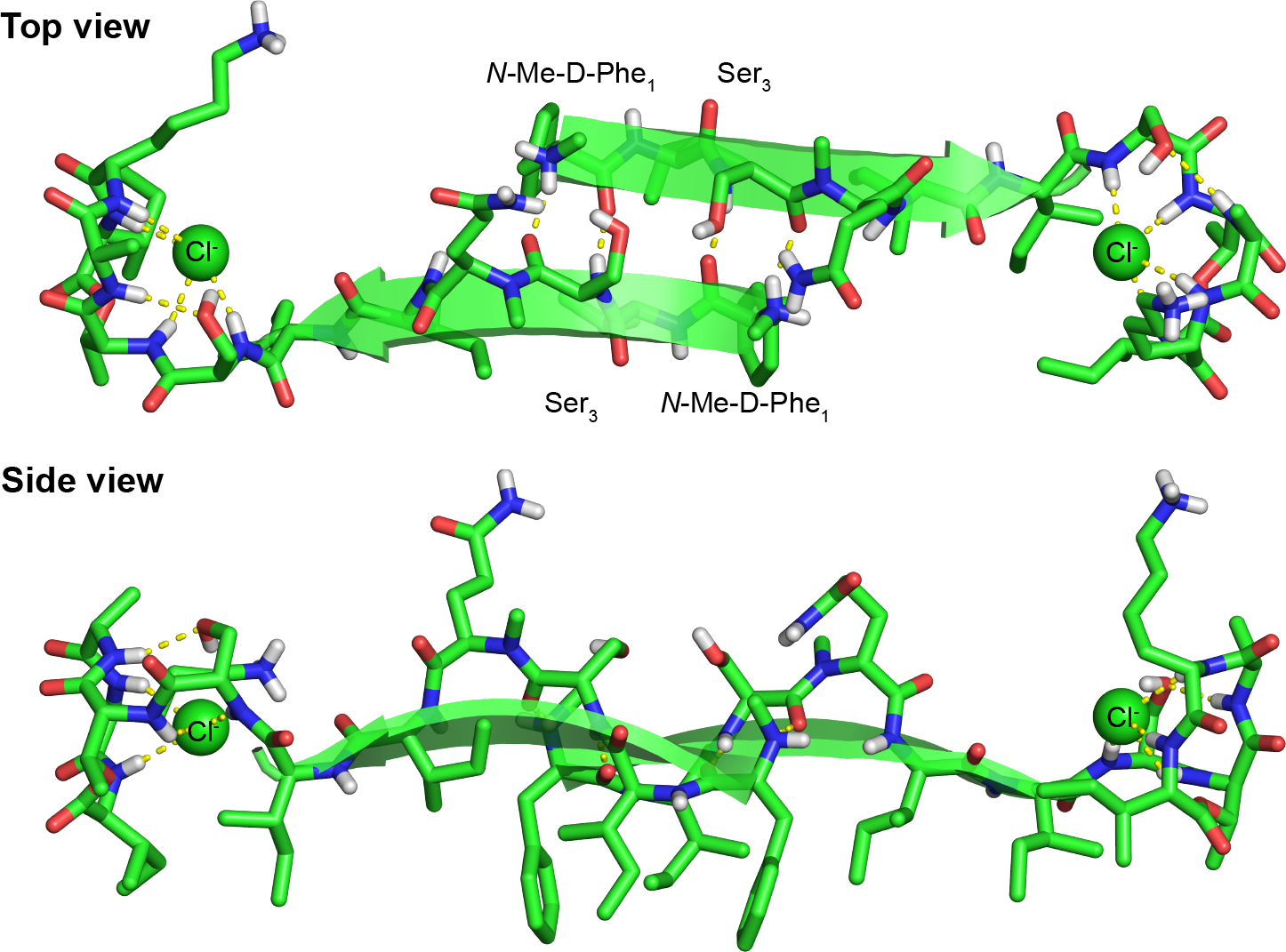
Hydrogen-bonded antiparallel β-sheet dimer in the crystal lattice of teixobactin analogue 1.

The dimers further assemble end-to-end in the crystal lattice, as illustrated in **Figure 4**. In this arrangement, the macrocycle of each molecule is in contact with the *N*-terminus of another molecule in the adjacent dimer, with the carbonyl group of Ala_9_ hydrogen bonding to the *N*-methylammonium group. Two macrocycles are in proximity at each interface between dimers, with the bound chloride anions ca. 7 Å apart. The end-to-end assembly of dimers is amphiphilic, and the hydrophobic surfaces pack together in the crystal lattice.

**Figure 4.**
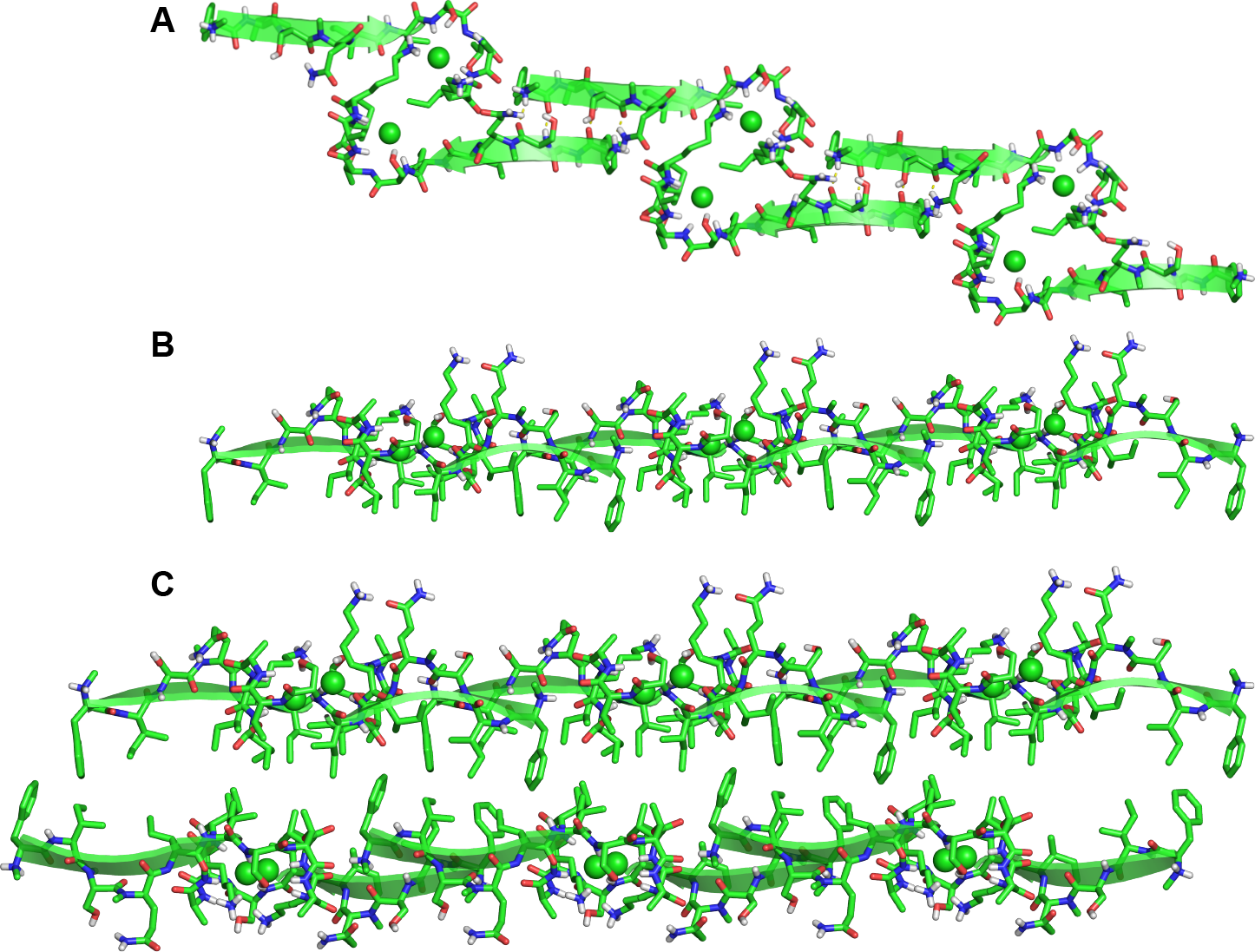
End-to-end dimer assemblies and their packing in the crystal lattice. (A) Top view. (B) Side view. (C) Hydrophobic packing in the crystal lattice.

## DISCUSSIONS

The dimers and other supramolecular interactions of *N*-Me-D-Gln_4_,Lys_10_-teixobactin (1) observed in the crystal state differ in several ways from those of *N*-methylated teixobactin analogues 2 and 3 (**Figure 5**). In the antiparallel β-sheet dimer of teixobactin analogue 1, residues 1–3 hydrogen bond, with *N*-Me-D-Phe_1_ pairing with Ser_3_. Analogue 2 also forms an antiparallel β-sheet dimer, but there is more overlap of the strands, with residues 1–6 forming the dimer interface (PDB 6E00). The dimers further assemble through antiparallel β-sheet interactions involving residues 3–7 of their outer edges. Analogue 3 forms an antiparallel β-sheet dimer, with residues 3–5 forming the dimer interface (PDB 6PSL). In the dimers formed by analogues 2 and 3, the *N*-terminus of one peptide molecule helps bind the anion that is grasped by the *C*-terminal region of the other molecule. In the dimers formed by analogue 1, however, only the C-terminal region participates in anion binding. The dimers of analogues 1 and 2 further assemble in the lattice to create amphiphilic sheets that pack through hydrophobic interactions. The dimers of analogue 3 further assemble through hydrophobic interactions to create three-fold symmetrical rods that extend through the crystal lattice.

**Figure 5.**
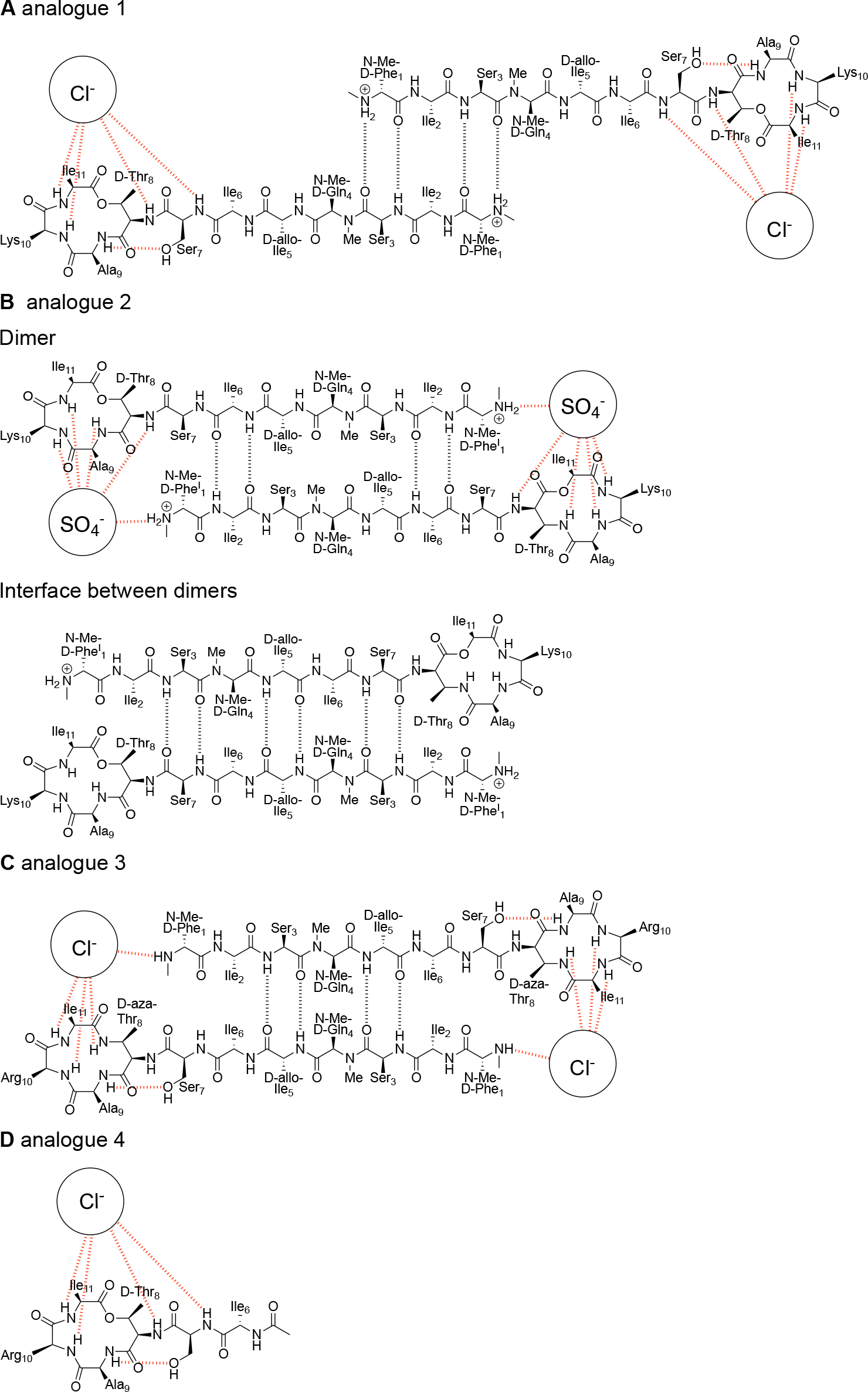
Chemical drawing of X-ray crystallographic structures of teixobactin analogues and their antiparallel β-sheet dimer interactions. (A) Analogue 1. (B) Analogue 2. (C) Analogue 3. (D) Analogue 4.

## CONCLUSIONS

Despite these differences, similarities exist across all of the X-ray crystallographic structures that we have determined. The analogues 1–3 each come together through antiparallel β-sheet interactions to create amphiphilic β-sheet dimers. The dimerization of these analogues likely reflects the propensity of teixobactin to form dimers and higher order assemblies that interact with bacterial cell membranes through hydrophobic interactions. Analogues 1–3 and truncated analogue 4 (CCDC 1523518) each bind anions in the crystal state, with analogues 1, 3, and 4 binding chloride and analogue 2 binding sulfate. The binding of these anions likely reflects the propensity of teixobactin to bind the pyrophosphate groups of lipid II and related cell wall precursors.

Supramolecular assembly is integral to the mechanism of action of teixobactin. *N*-Methylation at D-Gln_4_ helps prevent uncontrolled aggregation, permits crystallization of teixobactin analogues, and allows observation of conformation and supramolecular interactions at atomic resolution. In the crystal state, the teixobactin analogues adopt an amphiphilic conformation and form antiparallel β-sheet dimers though hydrogen bonding between the *N*-terminal tails. The macrocycles formed by residues 8–11 bind anions through hydrogen bonding. The hydrophobic surfaces of the dimers further pack through hydrophobic interactions. These observations further support a model in which teixobactin binds to Gram-positive bacteria through interactions of its hydrophobic side chains with the bacterial cell membrane, undergoes supramolecular assembly through β-sheet interactions, and binds the pyrophosphate groups of lipid II and related cell-wall precursors.

## EXPERIMENTAL SECTION

### General information

Methylene chloride (CH_2_Cl_2_) was passed through alumina under argon prior to use. Amine-free *N*,*N*-dimethylformamide (DMF) was purchased from Alfa Aesar. Fmoc-D-*allo*-Ile-OH was purchased from Santa Cruz Biotechnology. Fmoc-*N*-Me-D-Gln(Trt)-OH was purchased from ChemPep. Other protected amino acids were purchased from CHEM-IMPEX. 2-(Diphenylphosphino)benzoic acid was purchased from Arctom chemicals. Preparative reverse-phase HPLC was performed on a Rainin Dynamax instrument equipped with an Agilent Zorbax SB-C18 column. Analytical reverse-phase HPLC was performed on an Agilent 1260 Infinity II instrument equipped with a Phenomonex Aeris PEPTIDE 2.6μ XB-C18 column. HPLC grade acetonitrile (MeCN) and deionized water (18 MΩ) containing 0.1% trifluoroacetic acid (TFA) were used as solvents for both preparative and analytical reverse-phase HPLC. Deionized water (18 MΩ) was obtained from a Barnstead NANOpure Diamond purification system or a ThermoScientific Barnstead GenPure Pro water purification system. Glass solid-phase peptide synthesis vessels with fritted disks and BioRad Polyprep columns were used for solid-phase peptide synthesis. Teixobactin analogues **2a–10** were prepared and studied as the trifluoroacetate salts.

### Synthesis and crystallization of *N*-Me-D-Gln_4_,Lys_10_-teixobactin (1)

We synthesized *N*-Me-D-Gln_4_,Lys_10_-teixobactin (1) as the trifluoroacetate (TFA) salt as described. ^7^ Crystallization was performed using standard protein crystallography techniques. Briefly, *N*-Me-D-Gln_4_,Lys_10_-teixobactin (1) was dissolved in 18 MΩ deionized H_2_O at a concentration of 20 mg/mL. Suitable crystallization conditions were identified by screening experiments in a 96-well plate format with the aid of using screening kits from Hampton Research (including PEG/Ion, Index, and Crystal Screen kits). Each well was loaded with 100 μL of solution from the kits. Hanging drops were set up using a TTP Labtech Mosquito liquid handling instrument, with three 150-nL drops per well, with 1:1, 1:2, and 2:1 ratios of well solution and peptide stock solution. Crystals grew in conditions containing chloride with PEG3350 as precipitant.

To optimize crystal growth, we tested various conditions that included PEG3350 and MgCl_2_. In this optimization process, we varied the concentrations of PEG3350 (from 15% to 30%) and MgCl_2_ (from 0.20 M to 0.38 M) across a 4 × 6 matrix within a Hampton VDX 24-well plate. Hanging drops for the optimization experiments were prepared on glass slides by combining 1 or 2 μL of the *N*-Me-D-Gln_4_,Lys_10_-teixobactin solution with 1 or 2 μL of the well solution, using ratios of 1:1, 2:1, and 1:2. Crystals were assessed for X-ray diffraction using a Rigaku Micromax-007 HF diffractometer equipped with a Cu anode.

### X-ray data collection and processing

Data collection was conducted using the BOS/B3 software at the Advanced Light Source (ALS) on beamline 8.2.2. We collected X-ray diffraction at the longest wavelength possible (2.0663 Å, 6000 eV) to allow the use of chloride K-edge X-ray absorption (2822 eV) for single anomalous diffraction (SAD) phasing. Three data sets were acquired from three different crystals. Two sets of 360 images were acquired per crystal, with 1.0° rotation intervals (equivalent to two complete 360° rotations). The data sets were processed using XDS^8^, and the resulting data sets were merged employing BLEND^9^. To determine the structure, we used SAD phasing, implementing the Hybrid Substructure Search (HySS)^10^ module from the Phenix suite^11^.The anomalous signals from the chloride ions were used in this process. Initial electron density maps were generated by incorporating the chloride coordinates as starting positions in Autosol^12^. We also collected a higher-resolution data set at 0.9997 Å wavelength to allow the use of the molecular replacement with the structure determined from the 2.0663 Å data. X-ray diffraction data collection and processing information is summarized in **Table S1**.

### X-ray structure solution and refinement

The structure was refined using Phenix.refine^13^, and Coot^14^ was used for model building. During refinement, all B-factors were refined isotropically, and coordinates for hydrogen atoms were generated geometrically. For the non-proteinogenic amino acids (*N*-Me-D-Gln_4_, D-Thr_8_, and D-*allo*-Ile_5_), bond lengths, angles, and torsion restraints were generated using eLBOW^15^ in the Phenix software. The resulting structure was successfully used for molecular replacement with the 0.9997 Å resolution data set, which was refined in a similar fashion. Pentaethylene glycol dimethyl ether was included in the refinement to fill electron density associated with PEG3350. Final refinement and validation statistics are shown in **Table S2** (PDB 8U78).

### HPLC conditions and MS results

Analytical RP-HPLC was performed on a C18 column with an elution gradient of 5-67% CH_3_CN + 0.1% TFA over 15 min.

*N-Me-D-Gln*_*4*_,*Lys*_*10*_*-teixobactin (****1***). MS (ESI-MS): m/z: [M+H]^+^ Calcd for 1230.7; found 1230.5.

## ASSOCIATED CONTENT

### Supporting Information

The Supporting Information is available free of charge on the ACS Publications website at DOI:.

Supplementary figures and tables: HPLC and MS characterization data, X-ray crystallographic statistics, for *N*-Me-D-Gln_4_,Lys_10_-teixobactin (**1)**; Crystallographic coordinates of *N*-Me-D-Gln_4_,Lys_10_-teixobactin (**1**) deposited into the Protein Data Bank (PDB) with code 8U78.

### Notes

The authors declare no competing financial interest.

## ACKNOWLEDGEMENTS

Acknowledgements This work was supported by the National Institutes of Health (Grant Nos. AI121548 and AI137258). H.Y. acknowledges Allergan for fellowship support. We thank the Berkeley Center for Structural Biology (BCSB) of the Advanced Light Source (ALS) for synchrotron data collection. The BCSB is supported in part by the Howard Hughes Medical Institute. The ALS is supported by DOE Office of Science User Facility under Contract No. DE-AC02-05CH11231.

## TOC GRAPHIC

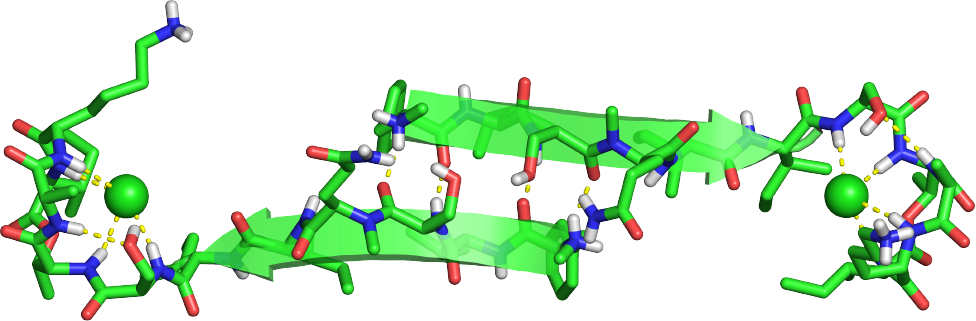

## Notes

### Competing Interest Statement

The authors have declared no competing interest.

### Summary of Updates

Updated manuscript and SI

