## Supplementary material for "Supramolecular Interactions of Teixobactin Analogues in the Crystal State": SI

### Supporting information:
### Supramolecular Interactions of Teixobactin Analogues in the Crystal State

Hyunjun Yang,[a] Adam G. Kreutzer,[a] and James S. Nowick[a,b,*]

[a] Department of Chemistry, University of California Irvine, Irvine, CA 92697.
[b] Department of Pharmaceutical Sciences, University of California Irvine, Irvine, CA 92697.


#### Table of Contents

**Supplementary Tables**

**Table S1.** X-ray crystallographic data collection and processing. S1

**Table S2.** X-ray crystallographic structure solution and refinement. S2

**Figure S1.** HPLC trace of *N*-Me-D-Gln$_4$,Lys$_{10}$-teixobactin. S3

**Figure S2.** Mass spectrum of *N*-Me-D-Gln$_4$,Lys$_{10}$-teixobactin. S4

#### Supplementary Tables

**Table S1.** X-ray crystallographic data collection and processing.

| | | |
|---|---|---|
| Diffraction source | ALS 8.2.2 beamline | ALS 8.2.2 beamline |
| Wavelength (Å) | 2.0663 | 0.9997 |
| Temperature (K) | 100 | 100 |
| Detector | 3×3 CCD array (ADSC Q315R) | 3×3 CCD array (ADSC Q315R) |
| Rotation range per image (°) | 1 | 1 |
| Total rotation range (°) | 720 | 720 |
| Space group | *P*43 | *P*43 |
| *a*, *b*, *c* (Å) | 25.86, 25.86, 109.50 | 24.88, 24.88, 106.32 |
| α, β, γ (°) | 90, 90, 90 | 90, 90, 90 |
| Resolution range (Å) | 2.022–27.40 | 1.148–24.88 |
| Total No. of reflections | 4147 | 20644 |
| Completeness (%) | 99.75 | 99.91 |
| $\langle I/\sigma(I)\rangle$ | 45.72 | 24.09 |
| Overall *B* factor from Wilson plot (Å$^2$) | 30.61 | 14.35 |

**Table S2.** Structure solution and refinement.

| | | |
|---|---|---|
| Resolution range (Å) | 27.38–2.40 (2.486–2.40) | 24.88–1.50 (1.554–1.50) |
| Completeness (%) | 99.75 (100) | 99.90 (100) |
| σ cutoff | $F > 1.36\sigma(F)$ | $F > 1.36\sigma(F)$ |
| No. of reflections, working set | 2805 (268) | 10318 (1055) |
| No. of reflections, test set | 282 (26) | 1030 (106) |
| Final $R_{cryst}$ | 0.2297 (0.2912) | 0.1544 (0.2188) |
| Final $R_{free}$ | 0.2494 (0.4508) | 0.1831 (0.2643) |
| Peptide | 8 | 8 |
| Water | 70 | 74 |
| Chloride | 9 | 14 |
| Ligand | | Pentaethylene glycol dimethyl ether |
| Bonds (Å) | 0.011 | 0.023 |
| Angles (°) | 1.44 | 2.30 |
| Average *B* factors (Å$^2$) | 30.61 | 20.41 |
| Ramachandran plot | | |
| Most favoured (%) | 87.5 | 100 |
| Allowed (%) | 12.5 | 0 |

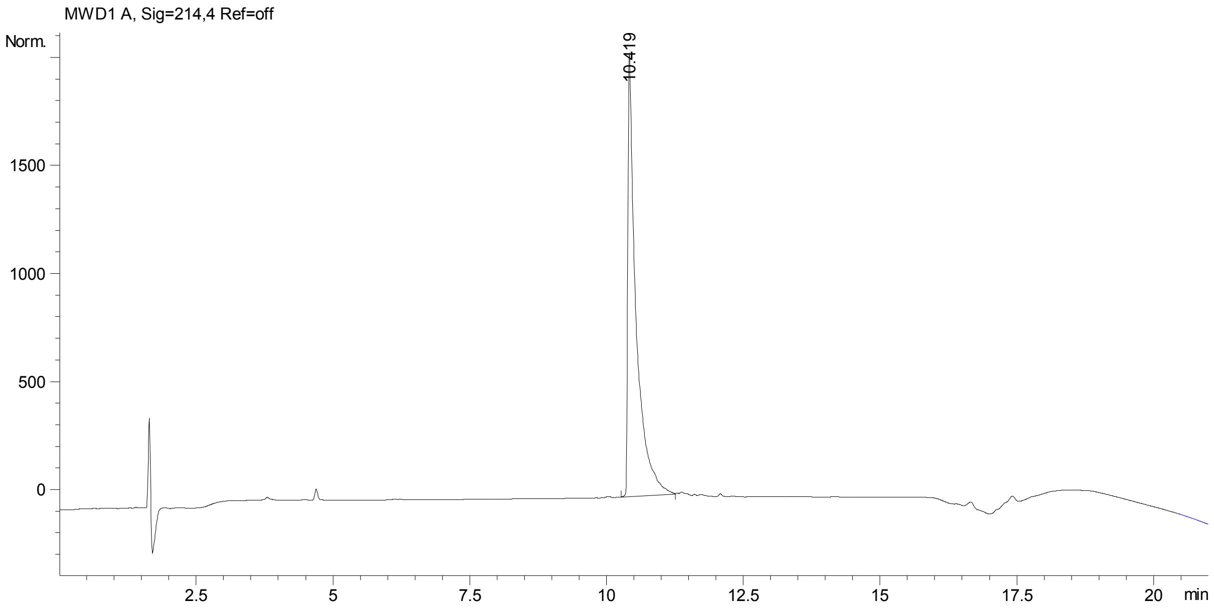


**Figure S1**. HPLC trace of *N*-Me-D-Gln4,Lys10-teixobactin.

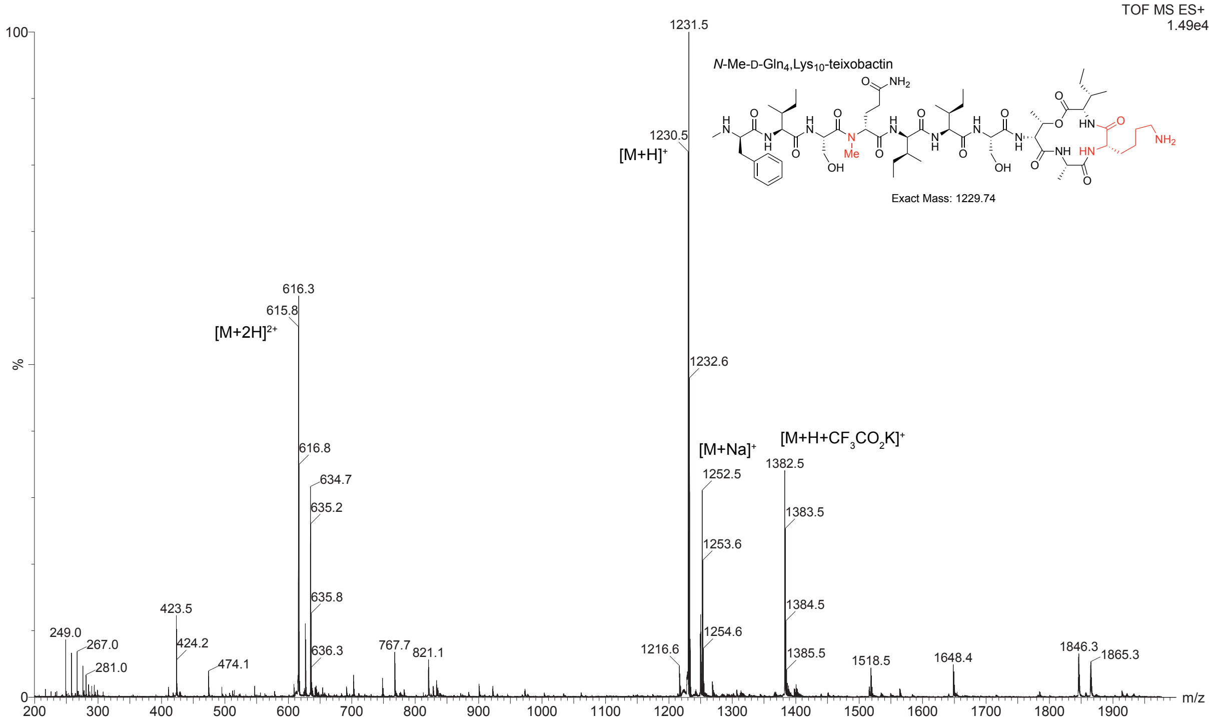


**Figure S2.** Mass spectrum of *N*-Me-D-Gln4,Lys10-teixobactin. Mass calculated for [M+H]+: 1230.74 and found 1230.5.
